# A mouse model for environmentally induced and reversible circadian arrhythmia using gradual exposure to a fragmented day-night cycle

**DOI:** 10.1101/2023.04.20.537697

**Authors:** Melissa E. S. Richardson, Chérie-Akilah Browne, Citlali I. Huerta Mazariegos

## Abstract

Arrhythmia is considered the most disrupted state of the biological circadian clock, and usually occurs when circadian regulatory genes are rendered non-functional, or the master clock (Suprachiasmatic Nucleus) is ablated. Since clock gene expression is aligned by the external solar day-night cycle to exhibit a 24-hour rhythm, we hypothesized that ill-timed light and dark exposure could negatively impact endogenous circadian clock function in mice. In this study, we present an environmentally driven approach to induce arrhythmia in mice that is also reversible. Using the previously characterized fragmented day-night cycle (FDN) where the 8-hour night is split into four 2-hour fragments and equally distributed across the 24-hour day, we show that mice gradually exposed to the FDN for 1 month lose their circadian rhythmicity. Furthermore, subsequent exposure to constant light or constant dark conditions does not yield typical circadian rhythms, but instead, reveals circadian arrhythmia. Finally, we show that the arrhythmic locomotion phenotype is reversible with one week of reintroduction to a 12 hr day-12 hr night cycle. This is the first study to show how the light-dark environment induces arrhythmia of an intact circadian clock and how it can be reversed.

## Introduction

Circadian rhythms control the daily reoccurrence of vital biological processes. The periodicity and alignment of circadian rhythms with the 24hr solar day are known as circadian photoentrainment and are important for the coordination of many biological processes [1, 2, 3, 4]. Aberrant light and dark exposure disrupt the natural day-night cycle and lead to vast circadian disruptions in mood, sleep, activity, metabolism, memory, and development [5, 6, 7, 8, 9]. The amount of time that each circadian-triggered event may last varies with the type of behavior in question. For instance, awakening and commencing daily activity occur almost simultaneously at a similar time each morning, however, the average waking time is short, 15-30 minutes, while most individuals remain active for a prolonged time, 16-18 hours [10]. However, whether circadian photoentrainment and the distribution of circadian-driven activity across the day may vary with sudden or gradual exposure to disruptive light cycles is less understood.

Circadian photoentrainment varies with the availability of light and dark [11, 12, 13]. Without light, mice exhibit an intrinsic circadian rhythm with an average period of 23.5 hours [14, 15, 16]. In contrast, constant 24-hour light exposure (LL) has been shown to induce a lengthened circadian rhythm with an average period of 25 hours (+/-0.5 hrs) [14, 17]. In addition to disrupted circadian photoentrainment, period lengthening is associated with many negative behavioral effects on development and mood [18, 19, 20]. A combination of light and dark has also been shown to induce period lengthening: fragmented day-night cycle (FDN), which divides an 8-hour night into 2-hour periods of darkness with intervening 4-hour light periods [21, 22]. The FDN cycle allows us to study how gradually changing the distribution of light and dark throughout the 24-hour day impacts circadian-driven behaviors such as the induction of period lengthening.

The goal of this study is to gradually fragment the 8-hr night of a 16hr light:8hr dark cycle, to determine the extent of night fragmentation necessary to disrupt circadian photoentrainment and induce period lengthening of the circadian rhythm using locomotor activity in mice. The study also explores the residual impact (known as aftereffects) of exposure to the FDN cycle on free-running rhythms in constant light and constant darkness (DD), which informs us of the potential impact of FDN exposure on circadian response to extreme conditions and maintenance of the intrinsic circadian period, respectively. The four major discoveries of this study are: 1) gradual exposure to the FDN cycle abolishes circadian rhythms compared to a sudden exposure to the same FDN cycle, 2) subsequent exposure to constant light unveils an arrhythmic clock, 3) exposure to constant darkness shows a weakened intrinsic clock rhythm, and finally, 4) the disruptive impact of FDN is reversed with 1-week exposure to a consolidated day-night cycle. This body of work broadens our understanding of how the timing and exposure modality to light and dark impact underlying circadian rhythms of locomotion.

## Results

### Circadian rhythms of locomotion are broadly distributed across the 24-hour day during gradual exposure to the fragmented day-night cycle (FDN-G) compared to sudden exposure (FDN-S)

Previous studies demonstrated that the fragmenting of the day-night cycle disrupted circadian photoentrainment, induced a lengthened period, increased activity during the light portion of the FDN cycle, and disrupted mood and development [21, 22]. Based on the previous findings, we hypothesized that there was a threshold for fragmenting the day-night cycle when mice would no longer exhibit circadian photoentrainment and switch to a lengthened circadian period (>25hrs). To determine the minimum level of fragmentation needed to disrupt circadian photoentrainment and induce period lengthening, the 4, 2-hour fragments of darkness in the FDN cycle were gradually separated by 1 hour each week until the traditional 4-hour light separation was accomplished for the FDN cycle (Figure 1A). Compared to the period lengthening observed following sudden exposure to FDN (FDN-S) (Figure 1B), mice gradually exposed to the FDN cycle (FDN-G) did not exhibit circadian period lengthening (Figures 1C, 1E, and 1H). Using the Lomb-Scargle periodogram to determine the periodicity, we found a strong peak at 24hrs in 16:8 LD mice and 25.4hrs in FDN-S mice, but no ∼24hr peak in FDN-G mice, indicating a lack of circadian rhythm (Figures 1E and 1H). Both FDN-S and FDN-G mice exhibit a non-circadian peak at 6hrs that mirrors the 4hr-light:2hr-dark cycle embedded within the FDN cycle (Figure 1E). FDN-G mice avoided light and restricted their activity to the dark fragments better than FDN-S mice similar to 16:8 LD mice (Figures 1D and 1G). There was no difference between the total daily activity level in 16:8 LD, FDN-S, and FDN-G conditions (Figure 1F). Compared to 16:8 LD and FDN-S, FDN-G mice exhibited wheel-running activity that was distributed across 20.33 hours ±0.36 (SEM) and was analyzed using the Circadian Index of Rhythmicity (CIR) scale, in which a value closer to 1 indicates a weak/no circadian rhythm (CIR scale also elaborated in the methods sections) (Figure 1I). The CIR value for 16:8 LD is 0.36 ±0.03 (SEM), FDN-S is 0.56 ±0.01 (SEM), and FDN-G is 0.85 ±0.01 (SEM) (Figure 1I). Together, these findings indicate that gradual exposure to the FDN cycle results in improved acute light suppression of locomotor activity and a non-circadian distribution of locomotor activity.

**Figure 1:**
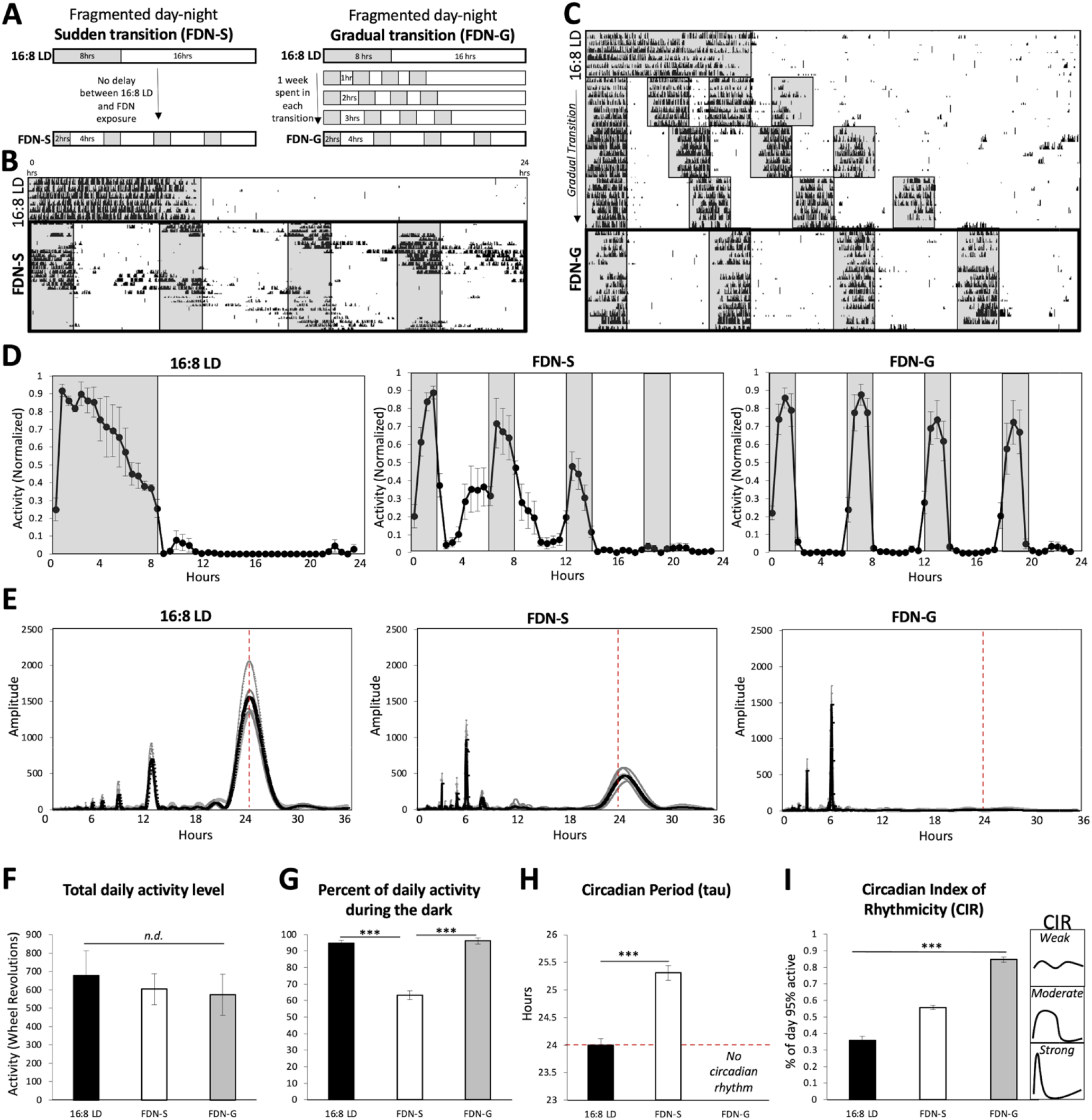
Circadian rhythms of locomotion are broadly distributed across the 24-hour day during gradual exposure to the fragmented day-night cycle (FDN-G) compared to sudden exposure (FDN-S). A) Schematic demonstrating the difference between the sudden exposure to the FDN cycle (FDN-S) and gradual exposure to the FDN cycle (FDN-G). B) Representative actograms of wheel-running activity during 16:8 LD to FDN-S and C) 16:8 LD to FDN-G. *n*=6 per group. D) Average of daily (across 7 days) wheel-running activity, collected in 30-minute bins, and normalized to the highest daily activity for each mouse for 16:8 LD, FDN-S, and FDN-G. n=6 per group. E) Lomb-Scargle periodograms across 7 days for 16:8 LD, FDN-S, and FDN-G; single-plotted (gray lines) with the average (black line). Alignment with the 24-hour day indicated by the dotted red line. *n*=6 per group, *F*= 106.68, *P*<0.0001***, peak average amplitudes: 1565.57, 473.99, Range: 702.93, 189.75, and ± SEM: 109.6, 29.5 (Comparison between all groups. Values for circadian (∼24hr) amplitude peaks for 16:8 LD and FDN-S are listed). F) Average total wheel-running activity across 7 days for 16:8 LD, FDN-S, and FDN-G. *n*=6 per group, *F*= 0.3161, *P*=0.7337, ranges: 850.9, 430.4, and 688.3, and ± SEM: 136.67, 83.69, 110.93, *n*.*d*.=no significant difference. G) Percent of total daily activity that occurred during the dark portion(s) for 16:8 LD, FDN-S, and FDN-G. *n*=6 per group, *F*= 53.57, *P*<0.0001***, ranges: 10.20, 28.4, 11.29, and ± SEM: 1.83, 2.57, 1.87. H) Circadian period (tau). Alignment with the 24-hour day indicated by the dotted red line for 16:8 LD and FDN-S. *n*=6 per group, difference between means (± SEM) 1.14 ± 0.14, *F*= 9.54, *P*<0.0001***, ranges: 0.3, 0.9, and ± SEM: 0.04, 0.13. I) Assessment of the period of circadian-driven wheel-running activity using the Circadian Index of Rhythmicity (CIR) scale for 16:8 LD, FDN-S, and FDN-G. Values closer to 0 indicate a strong rhythm while values close to 1 indicate weak circadian rhythms. *n*=6 per group, *F*= 0.40, *P*<0.0001***, and ranges: 3.50, 2.50, 2.50. ± SEM: 0.03, 0.01, 0.01.

### Arrhythmicity of circadian rhythm occurs during exposure to constant light (LL) following FDN-G

In addition to FDN-S, period lengthening is strongly associated with constant light conditions, which is considered highly disruptive to circadian photoentrainment and circadian-driven behaviors ([14, 18, 20] and Figure 2A). We hypothesized that since circadian period lengthening is observed in FDN-S, FDN-S mice would continue to lengthen their circadian period when exposed to LL, considering that previous studies showed that LL and FDN-S had similar behavioral phenotypes [21]. We also hypothesized that if FDN-G mice maintain any circadian rhythms, LL would induce period lengthening in FDN-G-exposed mice. Mice exposed to LL following FDN-S exhibited a continuous and extended period lengthening rhythm from FDN-S to LL, with a circadian period similar to mice previously exposed to 16:8 LD (Figure 2B, 2E, and 2G). In contrast to the period-lengthening phenotype in FDN-S, mice exposed to LL following FDN-G became arrhythmic, exhibiting no distinguishable peak on the activity profile or Lomb-Scargle periodogram (Figures 2C, 2D, 2E, and 2G). There was a reduction in wheel-running activity during LL following 16:8 LD, FDN-S, and FDN-G conditions, with previously FDN-G-exposed mice exhibiting the largest reduction (*P*=0.036) in activity (Figure 2F). Strong-moderate CIR scores represent the daily consolidation of circadian-driven locomotion observed in LL following 16:8 LD and FDN-S (Figures 2D and 2H). Circadian rhythms were abolished in LL for FDN-G exposed mice which exhibited a CIR score of 1.00 ±0.01 (SEM), which indicates that most of the locomotor activity was distributed across 24hrs instead of being restricted to 12hrs and 8.5hrs in LL-exposed 16:8 LD and FDN-S mice, respectively (Figures 2D and 2H). These findings indicate that gradual exposure to the FDN environment leads to circadian arrhythmia during constant light where a circadian rhythm of ∼25 hours is usually maintained.

**Figure 2:**
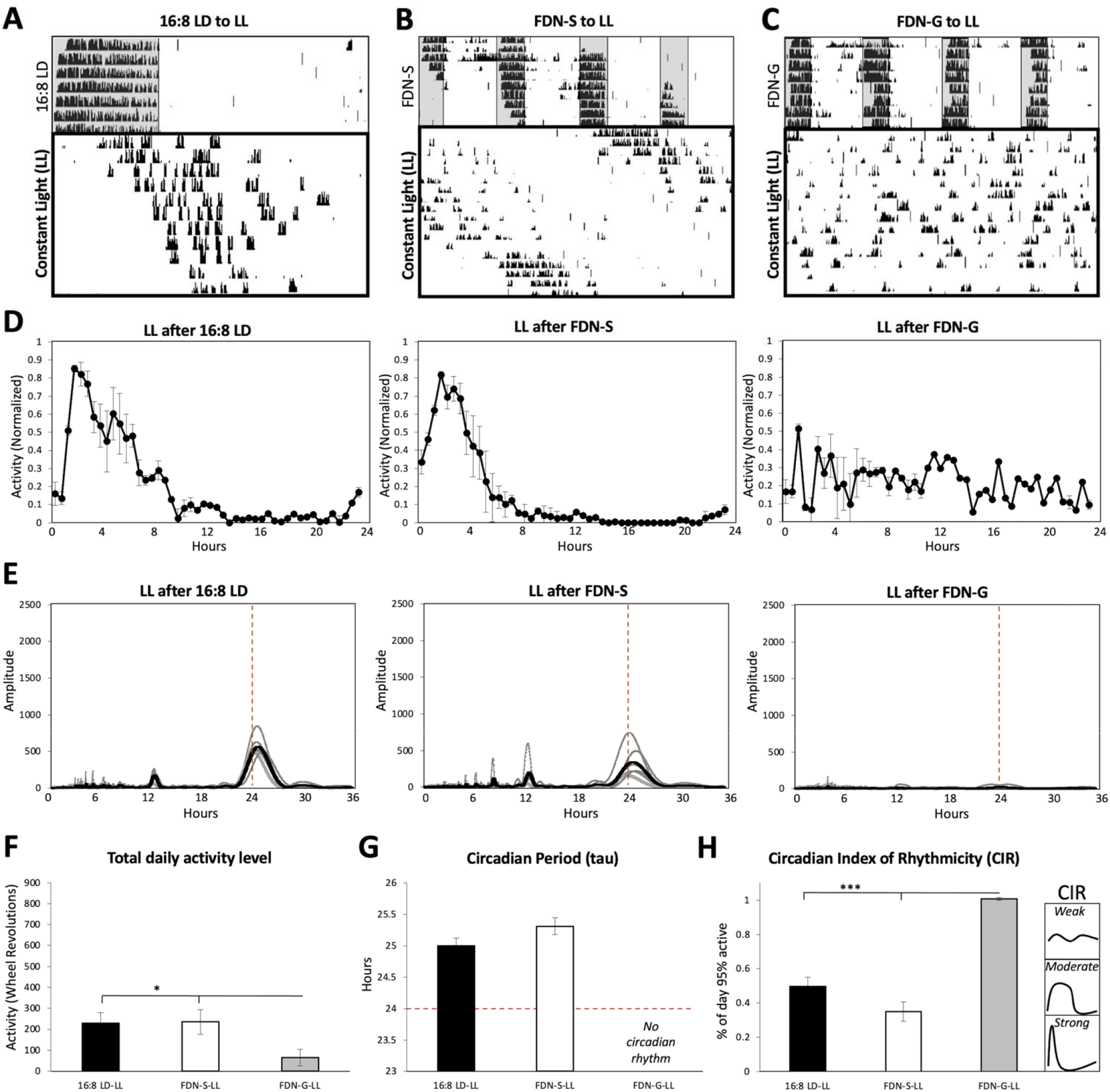
Arrhythmicity of circadian rhythm occurs during exposure to constant light (LL) following FDN-G. Representative actograms of wheel-running activity during constant light following A) 16:8 LD, B) FDN-S, and C) FDN-G. *n*=6 per group. D) Average of daily (across 7 days) wheel-running activity, collected in 30-minute bins, and normalized to the highest daily activity for each mouse. E) Lomb-Scargle periodograms across 7 days for LL following 16:8 LD, FDN-S, and FDN-G; single-plotted (gray lines) with the average (black line). Alignment with the 24-hour day indicated by the dotted red line. n=6 per group, F= 4.34, P=0.06, peak average amplitudes: 578.47, 354.71, Range: 362.95, 586.7, and ± SEM: 56.91, 91.05 (Comparison between 16:8 LD-LL and FDN-S-LL circadian (∼24hr) amplitude peak). F) Average total wheel-running activity across 7 days. *n*=6 per group, *F*= 4.16, *P*=0.036*, ranges: 308.0, 341.1, 230.1, and ± SEM: 42.86, 59.45, 35.02. G) Circadian period (tau). 24 hours is highlighted with a dotted red line for comparison. *n*=6 per group, Difference between means (± SEM) 0.30 ± 0.20, *F*= 1.29, *P*=0.17, ranges: 1.0, 0.9, and ± SEM: 0.15, 0.14. H) Assessment of the period of circadian-driven wheel-running activity using the Circadian Index of Rhythmicity (CIR) scale. Values closer to 0 indicate a strong rhythm while values close to 1 indicate weak circadian rhythms. *n*=6 per group, *F*= 43.75, *P*<0.0001***, and ranges: 11.00, 9.00, 1.00. ± SEM: 1.49, 1.37, 0.20.

### Intrinsic circadian rhythm is temporarily arrhythmic during constant dark exposure following FDN-G

Analysis of the intrinsic circadian rhythm, which occurs in the absence of light stimuli, informs us of the internal rhythmic state of the clock, which typically exhibits a strong circadian period of approximately 23.5 hours in mice ([14, 15, 16] and Figures 3A, 3D, 3E, and 3G). To determine if the FDN-S and FDN-G cycles were detrimental to the intrinsic clock function, we removed light stimuli following FDN-S and FDN-G and observed the activity rhythms (Figures 3B and 3C). The level of wheel-running activity was similar in DD for all experimental groups (Figure 3F). During DD but following FDN-S and 16:8 LD, mice immediately switched to a shortened circadian period and exhibited strong amplitude peaks on the Lomb-Scargle periodogram (Figures 3A, 3B 3E, and 3G). However, when exposed to DD for 12 days, FDN-G mice exhibit circadian arrhythmia for the first 5 days, as quantified by a reduced amplitude (F= 9.52, P<0.0001) on the Lomb-Scargle periodogram and a very weak rhythm with a CIR score of 0.90 ±0.69 (SEM) (*F*= 48.65, *P*<0.0001) (Figures 3E and 3H). However, during the latter portion of the DD exposure period (days 6-12), the previously exposed FDN-G mice recovered a normal shortened circadian rhythm of 23.65 hours ±0.04 (SEM), increased amplitude similar to FDN-S and 16:8 LD mice and exhibited a strong circadian rhythm with a CIR score of 0.53 ±0.79 (SEM) (Figures 3C, 3D, 3E, 3G, and 3H). Together, these findings confirm that gradual exposure to the FDN cycle results in temporary circadian arrhythmia in DD, which lasted for the first five days of the total 12 days in DD.

**Figure 3:**
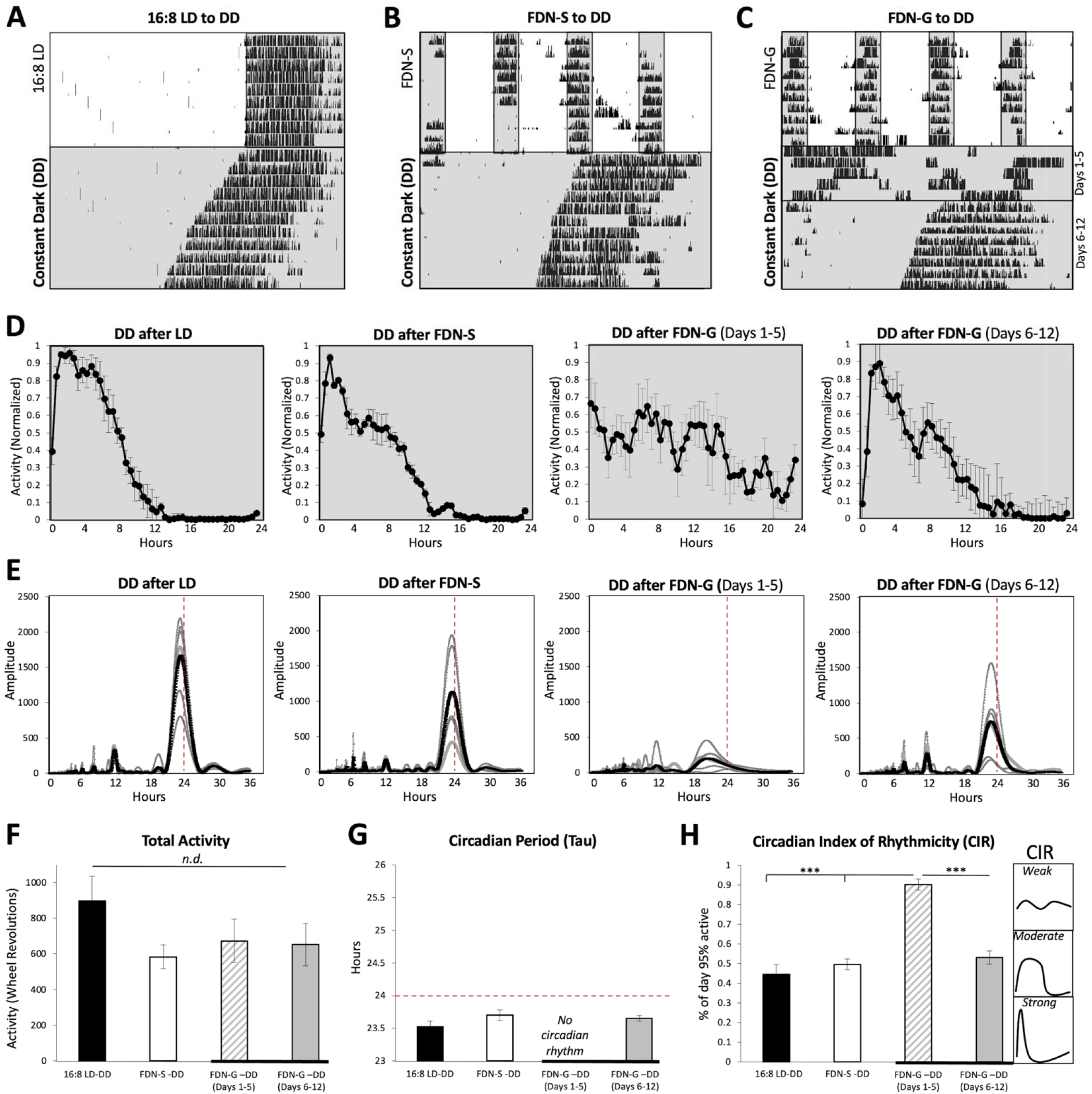
FDN-G exposure results in arrhythmia of the intrinsic circadian rhythm during constant darkness (DD). Representative actograms of wheel-running activity during constant darkness following A) 16:8 LD, B) FDN-S, and C) FDN-G. *n*=6 per group. D) Average of daily (across 7 days) wheel-running activity, collected in 30-minute bins, and normalized to the highest daily activity for each mouse. E) Lomb-Scargle periodograms across 7 days for DD following 16:8 LD, FDN-S, and FDN-G; single-plotted (gray lines) with the average (black line). Alignment with the 24-hour day indicated by the dotted red line. n=6 per group, F= 9.52, P<0.0001***, peak average amplitudes: 1671.82, 1132.36, 212.72, 740.94, Range: 1390.03, 1525.79, 399.08, 1369.8 and ± SEM: 228.28, 248.87, 57.56, 206.14 (Comparison between 16:8 LD-DD, FDN-S-DD, and FDN-G-DD circadian (∼24hr) amplitude peak). F) Average total wheel-running activity across 7 days. *n*=6 per group, *F*= 1.33, *P*=0.29, ranges: 867.4, 583.5, 739.9, 715.4 and ± SEM: 141.9, 76.94, 121.9, 118.4. G) Circadian period (tau). 24 hours is highlighted with a dotted red line for comparison. *n*=6 per group, *F*= 2.65, *P*=0.10, ranges: 0.40, 0.40, 0.30, and ± SEM: 0.05, 0.06, 0.04. H) Assessment of the period of circadian-driven wheel-running activity using the Circadian Index of Rhythmicity (CIR) scale. Values closer to 0 indicate a strong rhythm while values close to 1 indicate weak circadian rhythms. *n*=6 per group, *F*= 48.65, *P*<0.0001***, and ranges: 4.50, 5.00, 5.00, 5.00. ± SEM: 0.66, 0.76, 0.69, 0.79.

### The effects of the FDN-G on locomotion are temporary. Normal rhythms recovered in 1 week

In a previous study, we showed that re-introduction to a 12:12 LD cycle following FDN-S reversed negative effects on mood in 1-2 weeks [21]. Therefore, we wanted to determine if exposure to 12:12 LD can recover circadian photoentrainment as well as the lengthened and shortened circadian rhythms associated with LL and DD, respectively, in FDN-G-exposed mice. FDN-G mice exposed to 1-week of 12:12 LD showed immediate recovery of an entrained 24-hour circadian rhythm (Figures 4A, 4B, & 4C - middle panel). When the same previously FDN-G exposed mice were suddenly re-exposed to the FDN cycle (FDN-S) following 12:12 LD, they demonstrated recovery of a circadian rhythm, increased amplitude, and exhibited the period-lengthening phenotype previously observed with the FDN-S mice (Figures 4A, 4D, 4E, 4F, and 4G). Similarly, exposure of FDN-G mice to LL and DD after 1 week in 12:12 LD yields a recovery of circadian rhythms, increased amplitudes, and circadian periods typical of LL and DD following an LD cycle (Figures 4B-4F). The recovered rhythms (CIR analysis) of the FDN-G mice were not different from the control mice that were only exposed to an LD cycle before FDN-S, LL, or DD (Figure 4G). These findings indicate that the disrupted circadian rhythm during FDN-G is reversible with 1 week of 12:12LD exposure.

**Figure 4:**
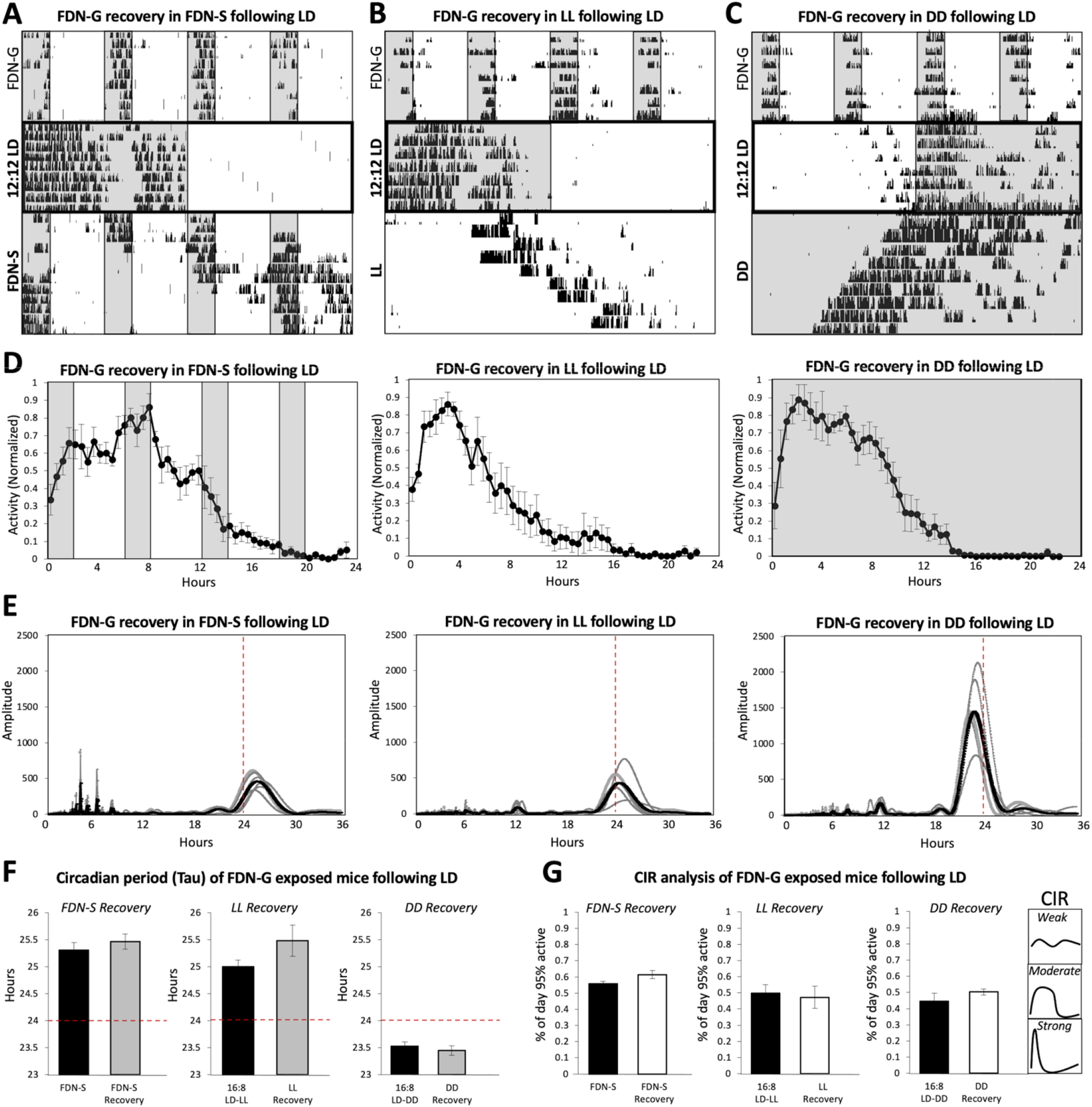
One week of consolidated light-dark exposure following FDN-G resets the disrupted circadian rhythms. A) Representative actograms of wheel-running activity during A) FDN-S, B) LL, and C) DD following FDN-G and 12:12 LD exposure. *n*=6 per group. D) Average of daily (across 7 days) wheel-running activity, collected in 30-minute bins, and normalized to the highest daily activity for each mouse. E) Lomb-Scargle periodograms across 7 days for FDN-S, LL, and DD following FDN-G and 12:12 LD exposure; single-plotted (gray lines) with the average (black line). Alignment with the 24-hour day indicated by the dotted red line. n=6 per group, F= 9.52, P<0.0001***, peak average amplitudes: 1671.82, 1132.36, 212.72, 740.94, Range: 1390.03, 1525.79, 399.08, 1369.8 and ± SEM: 228.28, 248.87, 57.56, 206.14 (Comparison between 16:8 LD-DD, FDN-S-DD, and FDN-G-DD circadian (∼24hr) amplitude peak). F) Circadian period (tau) during FDN-S (*n*=6 per group, *F*= 1.23, *P*=0.15, range: 0.9, 0.8, and ± SEM: 0.13, 0.14), LL (*n*=6 per group, *F*= 3.57, *P*=0.18, range: 1.0, 2.0, and ± SEM: 0.15, 0.29), and DD (*n*=6 per group, *F*= 2.95, *P*=0.47, range: 0.4, 0.7, and ± SEM: 0.05, 0.09) compared to mice previously exposed to 16:8 LD only. 24 hours is highlighted with a dotted red line for comparison. G) Assessment of the period of circadian-driven wheel-running activity using the Circadian Index of Rhythmicity (CIR) scale during FDN-S (*n*=6, *F*= 2.67, *P*=0.18, range: 2.50, 3.50, and ± SEM: 0.35, 0.58), LL (*n*=6, *F*=1.24, *P*=0.79, range: 11.00, 10.50 and ± SEM: 1.47, 1.64), and DD (*n*=6, *F*= 2.13, *P*=0.07, range: 4.50, 2.50, and ± SEM: 0.66, 0.45) compared to mice previously exposed to 16:8 LD only. Values closer to 0 indicate a strong rhythm while values close to 1 indicate weak circadian rhythms.

## Discussion

With frequent exposure to light at night, daytime napping, and shift-work schedules, it is often difficult to maintain circadian photoentrainment. Our study demonstrates how light and dark exposure can impact three levels of circadian rhythm maintenance: 1) circadian photoentrainment, the most ideal for physiological homeostasis [4], 2) circadian period lengthening, a disruptive deviation from circadian entrainment but still a positive indicator of a functional clock that is maintained under a number of disruptive T-cycles [23, 24, 25, 26, 27], and 3) circadian arrhythmia, the most disruptive state of the circadian clock that is usually associated with SCN (suprachiasmatic nucleus) ablation or mice with mutant clock gene components such as BMAL1 [28, 29, 30]). Our FDN-G paradigm led to circadian arrhythmia in locomotor activity. However, unlike clock mutants, the effects of FDN-G were not permanent or long-term, as the circadian rhythm was reset after 1-week of exposure to 12:12 LD and 5 days in DD. Therefore, we propose that the FDN-G cycle could be used as a model for temporary arrhythmia induction in mice.

There have been numerous studies on disrupting circadian photoentrainment with light, however, inducing circadian arrhythmia using changes to the 24hr solar day has proven more elusive, although one study used extreme exposure to constant light (130 days) to eventually induce circadian arrhythmia [11, 13, 31, 32]. We are interested in light-dark disruptions that are socially relevant to the 24hr solar day. Some studies used non-24-hr T-cycles with two separate light-dark periods (LDLD cycles), referred to as bifurcation, which disrupts circadian photoentrainment and results in the alignment of activity rhythms to the two periods of darkness and does not induce circadian arrhythmia when exposed to constant dark conditions [13, 31, 33]. Although the achievement of bifurcation is regarded as advantageous for the adaptability of activity rhythms to extreme changes to the 24hr solar cycle (jet lag, shiftwork, space travel), considering bifurcation as a therapeutic option for daily adaptability requires the context of the solar day [34, 35]. Additionally, authors studying bifurcation including Gorman and Elliot, reported that sudden or gradual exposure to T18 or T30 yielded no difference in the ability of animals to achieve bifurcated rhythms [12, 13, 31, 33, 34]. These studies on bifurcation are different from the current study which shows how a gradual, not sudden, exposure to 4 LD cycles within 24 hours (LDLDLDLD: quadfurcation), leads to circadian arrhythmia in constant dark conditions. Taken together with studies on LD bifurcation, we uncover a threshold for disruptive LD fragmentation, where LD quadfurcation (FDN-G) leads to circadian arrhythmia.

The timing of exposure to light and dark strongly influences the circadian clock and behavior [3, 36]. In our fragmented day-night paradigm, FDN-G mice aligned their activity with the dark fragments and avoided the light exquisitely, similar to the LDLD bifurcation paradigm, but in contrast, lost circadian rhythmicity in constant conditions [12, 13, 33, 34]. Additionally, FDN-G mice instantly regained circadian rhythmicity following FDN-G upon exposure to the 1-week 12:12 LD cycle. FDN-G alignment with the dark fragments was not a circadian response, but likely another mechanism, since a similar finding was observed in BMAL1 and Per2/Cry1 mutants, which show similar alignment to LD cycles but arrhythmia in DD and LL [28, 37]. This means that light can impact behavior independently of a functional biological clock and has previously been described as an acute effect of light on behavior that overrides the influence of the circadian clock, known as negative masking [38, 39]. Therefore, the typical confinement of activity to the dark portion of the day is not completely under the control of the circadian clock, but rather a combination of circadian and acute light response drivers.

Researchers often employ wheel-running activity and the ClockLab software to record and analyze circadian rhythms in mice (14, 40, 41, 42, 43]. Using data exported from ClockLab to increase the analytic resolution of our study, we examined multiple facets of circadian locomotion rhythms such as periodicity, amplitude, consistency of activity between animals, and the distribution of activity across the day. The overall presence of a circadian rhythm and relative confidence of the algorithm (amplitude) were measured by a built-in analysis tool in ClockLab, the Lomb-Scargle periodogram, which is better suited for shorter periods of rhythm assessment (our data was 5-7 days on average) as compared to the Chi-square periodogram [44, 45]. Additionally, we devised the Circadian Index of Rhythmicity (also referred to as the CIR scale), which directly describes the dispersion of locomotor behavior across the day, by determining how many hours of the day it takes to complete 95% of the total activity, with the highest value (1) indicating maximum dispersed activity and thus, a lack of circadian behavior. The CIR scale could be applied to a range of circadian-driven behaviors to determine the quantifiable fraction of the day different circadian behavior typically occupy, which will make the assessment of deviating behaviors easier to understand.

## Conclusion

Adaptation to a disruptive LD environment by confining activity to the available dark period may seem like a positive behavior for mice. However, here we show that is completely the opposite, as gradual adaptation, and confinement to the four periods of darkness in the FDN-G paradigm leads to circadian arrhythmia. The state of the circadian clock as it relates to the external environment is central to maintaining the synchronous internal balance necessary to support homeostatic mechanisms of a broad range of biological functions. Future studies would be necessary to determine the best timing for light and dark exposure in humans, which is important to support daily rhythmic coordination in our bodies as well as recuperative health for the ill who are often exposed to irregular light schedules in hospitals and care facilities.

## Methods

### Mice

Age-matched C57BL/6NCrl males between 5 and 7 months old, were used for our experiments. Male mice were used in this study due to findings of increased variability in circadian-driven behaviors such as locomotion due to hormone variability associated with the 4-5 estrous cycle in female mice [2, 46]. Future studies that can account for estrous-related variability will be considered in gendered circadian studies. Mice were housed and treated in accordance with NIH, ARRIVE, and IACUC guidelines and used protocols approved by the Oakwood University Animal Care and Use Committee (OUACUC). Six mice were used per experimental group.

### Light cycles

#### Consolidated night (12:12 LD) cycle

12:12 LD is one of the most used light cycles to maintain and breed mice [47]. It consists of 12 hours of darkness and 12 hours of light that repeats every 24 hours. Mice were bred and maintained in 12:12 LD before experiments. The light intensity used in our experiments was ∼500 lx of white light (similar to office lighting).

#### Consolidated night (16:8 LD) cycle

Mice were transferred to the 16:8 LD cycle at least 2 weeks before any experimental changes in the light-dark environment occurred.

#### Fragmented day-night-sudden (FDN-S) cycle

The fragmented day-night cycle used in this study consists of 4-alternating cycles of 2 hours of darkness and 4 hours of light each 24-hour day (also called T6 cycle in [21]). Mice were transferred immediately from 16:8 LD to FDN to create the FDN-S condition.

#### FDN-G (gradual) cycle

Gradual exposure to the FDN cycle started with fragmenting the 8-hour night into 2-hour fragments, separated by 1 hour of light for 1 week. Each subsequent week, the amount of light between fragments was increased by an hour to 2 hours, 3 hours, and eventually, the 4-hour light exposure experience by the traditional FDN-S cycle explained above. Mice were maintained for 7-10 days in FDN-G before exposure to LL, DD, or LD paradigms (figures 2-4).

### Circadian assessment of locomotion using wheel-running activity

Mice (male) were placed in cages with a 6-inch Actimetrics Wireless Low-profile Running Wheel. Analyses and monitoring of wheel-running activity were conducted with ClockLab (Actimetrics). Analyses include total activity, period length, and activity distribution across the light and dark periods of the day. Raw data for each animal were exported for statistical analysis. The Lomb-Scargle periodogram was used to analyze amplitude and periodicity (tau). The activity onset for each free-running animal (LL and DD) was aligned to circadian time 12 using the Tau function (which matched the period calculation by the Lomb-Scargle periodogram) in the Actogram window of Clocklab, before exporting data for analysis and creating activity distribution graphs.”

### Circadian Index of Rhythmicity (CIR)

Circadian distribution of activity across the 24-hour day was determined by calculating the average wheel revolution in 30-minute bins over 7 days for each mouse. The onset of activity (circadian time 12) was aligned with bin 1 (the first 30 minutes of analysis). If the circadian period was longer or shorter than 24 hours, the tau was first adjusted to have the same onset time for activity each day before exporting daily averages. CIR= Time (in sums of 30 mins bins) to accomplish 95% of daily locomotion/24. CIR scores of 0.33 and less are considered a strong circadian rhythm. CIR scores 0.34-0.66 are considered a moderate circadian rhythm. CIR scores of 0.67 or more are considered a weak-absent circadian rhythm.

### Statistical analyses

Statistical analyses were conducted with Prism GraphPad, version 9.4.1. Six mice were used for each experiment; mice were not reused in different experiments. Where a comparison of three conditions was conducted, the one-way-ANOVA analysis was performed with a Tukey’s multiple comparisons post-test, because each data set was compared to the other two data sets. Where a comparison of two groups was conducted, the unpaired t-test, two-tailed was used. The alpha for all tests is 0.05. The distribution of data sets exhibited a Gaussian distribution of normality. The error bars represent the standard error of the mean. Descriptive statistics including the range, p-values, SEM, and F-values are included in each figure legend.

## Data availability

All data are available on reasonable request, directed to the corresponding author, Melissa E S Richardson.

## Acknowledgments

Thanks to Mr. Shiraz Davis and Dr. Vanterpool of the Department of Biological Sciences at Oakwood University for their staff and administrative support.

## Author contributions

**MESR** conceptualized the research questions, conducted the behavioral experiments, analyzed the data, created the figures, co-wrote all drafts, and edited all drafts. **CAB** analyzed the data, co-wrote all drafts, and edited all drafts. **CIHM** conducted the behavioral experiments, analyzed the data, and edited drafts.

## Additional Information

### Ethics declarations

This study aligns with the ARRIVE guidelines, uses enough mice for experiments, avoids invasive methods, and uses ethical practices.

## References

1. Blume, C., Garbazza, C., & Spitschan, M. Effects of light on human circadian rhythms, sleep and mood. Somnologie. 23(3), 147–156 (2019).

2. Jud, C., Schmutz, I., Hampp, G., Oster, H., & Albrecht, U. A guideline for analyzing circadian wheel-running behavior in rodents under different lighting conditions. Biol. Proced. Online. 7(1), 101–116 (2005).

3. Foster, R. G., Hughes, S., & Peirson, S. N. Circadian photoentrainment in mice and humans. Biology. 9(7), 180 (2020).

4. Bruce, V. G. Environmental entrainment of circadian rhythms. Cold Spring Harb. Symp. Quant. Biol. 25, 29–48 (1960).

5. Heyde, I., & Oster, H. Differentiating external zeitgeber impact on peripheral circadian clock resetting. Sci. Rep, 9(1), 1–13 (2019).

6. Hasan, S., Foster, R. G., Vyazovskiy, V. V., & Peirson, S. N. Effects of circadian misalignment on sleep in mice. Sci. Rep. 8(1), 1–13 (2018).

7. Alves-Simoes, M., Coleman, G., & Canal, M. M. Effects of type of light on mouse circadian behaviour and stress levels. Lab. Anim. 50(1), 21–29 (2016).

8. Schilperoort, M., et al. Disruption of circadian rhythm by alternating lightLJdark cycles aggravates atherosclerosis development in APOE* 3LJLeiden. CETP mice. J. Pineal Res. 68(1), e12614 (2020).

9. Ameen, R. W., Warshawski, A., Fu, L., & Antle, M. C. Early life circadian rhythm disruption in mice alters brain and behavior in adulthood. Sci. Rep. 12(1), 1–13 (2022).

10. Roenneberg, T., Wirz-Justice, A., & Merrow, M. Life between clocks: daily temporal patterns of human chronotypes. J. Biol. Rhythms. 18(1), 80–90 (2003).

11. Gorman, M.R. Exotic photoperiods induce and entrain split circadian activity rhythms in hamsters. J Comp Physiol A. 187, 793–800 (2001).

12. Rosenthal, S. L., Vakili, M. M., Evans, J. A., Elliott, J. A., & Gorman, M. R. Influence of photoperiod and running wheel access on the entrainment of split circadian rhythms in hamsters. BMC neurosci. 6, 1–13 (2005).

13. Sun, J., Joye, D. A., Farkas, A. H., & Gorman, M. R. Photoperiodic requirements for induction and maintenance of rhythm bifurcation and extraordinary entrainment in male mice. Clocks & sleep. 1(3), 290–305 (2019).

14. EckelLJMahan, K., & SassoneLJCorsi, P. Phenotyping circadian rhythms in mice. Curr. Proc. 5(3), 271–281 (2015).

15. Hughes, A.T.L., et al. Timed daily exercise remodels circadian rhythms in mice. Commun. Biol. 4, 761 (2021).

16. Aton, S. J., Block, G. D., Tei, H., Yamazaki, S., & Herzog, E. D. Plasticity of circadian behavior and the suprachiasmatic nucleus following exposure to non-24-hour light cycles. J. Biol. Rhythms. 19(3), 198–207 (2004).

17. Hughes, A. T., et al. Constant light enhances synchrony among circadian clock cells and promotes behavioral rhythms in VPAC2-signaling deficient mice. Sci. Rep. 5(1), 1–12 (2015).

18. Ohta, H., Mitchell, A. & McMahon, D. Constant Light Disrupts the Developing Mouse Biological Clock. Pediatr. Res. 60, 304–308 (2006).

19. Oliver, P. L., et al. Disrupted circadian rhythms in a mouse model of schizophrenia. Curr. Biol. 22(4), 314–319 (2012).

20. Tapia-Osorio, A., Salgado-Delgado, R., Angeles-Castellanos, M., & Escobar, C. Disruption of circadian rhythms due to chronic constant light leads to depressive and anxiety-like behaviors in the rat. Behav. Brain Res. 252, 1–9 (2013).

21. Richardson, M. E., Brown, D., Honore, D., & Labossiere, A. Fragmented day-night cycle induces period lengthening, lowered anxiety, and anhedonia in male mice. Behav. Brain Res. 413, 113453; 10.1016/j.bbr.2021.113453 (2021).

22. Richardson, M. E., Boutrin, M. C., Chunn, S., & Hall, M. Fragmented day-night cycle induces reduced light avoidance, excessive weight gain during early development, and binge-like eating during adulthood in mice. Physiol. Behav. 113851; 10.1016/j.physbeh.2022.113851 (2022).

23. Chen, R., Seo, D. O., Bell, E., Von Gall, C., & Lee, C. Strong resetting of the mammalian clock by constant light followed by constant darkness. J. Neurosci. 28(46), 11839–11847 (2008).

24. Park, N., Cheon, S., Son, G. H., Cho, S., & Kim, K. Chronic circadian disturbance by a shortened light-dark cycle increases mortality. Neurobiol. Aging. 33(6), 1122–e11 (2012).

25. Altimus, C. M., et al. Rods-cones and melanopsin detect light and dark to modulate sleep independent of image formation. PNAS. 105(50), 19998–20003 (2008).

26. LeGates, T. A., et al. Aberrant light directly impairs mood and learning through melanopsin-expressing neurons. Nature. 491(7425), 594–598 (2012).

27. Ono, D., Honma, S., & Honma, K. I. Postnatal constant light compensates Cryptochrome1 and 2 double deficiency for disruption of circadian behavioral rhythms in mice under constant dark. PloS one. 8(11), e80615 (2013).

28. Izumo, M., et al. Differential effects of light and feeding on circadian organization of peripheral clocks in a forebrain Bmal1 mutant. Elife. 3, 04617; 10.7554/eLife.04617 (2014).

29. Abe, Y. O., et al. Rhythmic transcription of Bmal1 stabilizes the circadian timekeeping system in mammals. Nat. commun. 13(1), 1–12 (2022).

30. Tso, C. F., et al. Astrocytes regulate daily rhythms in the suprachiasmatic nucleus and behavior. Curr. Biol. 27(7), 1055–1061 (2017).

31. Gorman, M. R., & Elliott, J. A. Focus: Clocks and Cycles: Exceptional Entrainment of Circadian Activity Rhythms With Manipulations of Rhythm Waveform in Male Syrian Hamsters. YJBM. 92(2), 187 (2019).

32. Ohta, H., Yamazaki, S., & McMahon, D. G. Constant light desynchronizes mammalian clock neurons. Nat. Neurosci. 8(3), 267–269 (2005).

33. Harrison, E., et al. Extraordinary behavioral entrainment following circadian rhythm bifurcation in mice. Sci Rep. 6, 38479 (2016).

34. Walbeek, T. J., Harrison, E. M., Soler, R. R., & Gorman, M. R. Enhanced circadian entrainment in mice and its utility under human shiftwork schedules. Clocks & sleep. 1(3), 394–413. (2019).

35. Noguchi, T., et al. Circadian rhythm bifurcation induces flexible phase resetting by reducing circadian amplitude. Eur. J. Neurosci. 51(12), 2329–2342 (2020).

36. Li, Y., & Androulakis, I. P. Light entrainment of the SCN circadian clock and implications for personalized alterations of corticosterone rhythms in shift work and jet lag. Sci. Rep. 11(1), 1–19 (2021).

37. Abraham, D., et al. Restoration of circadian rhythmicity in circadian clock-deficient mice in constant light. J. Biol. Rhythms. 21(3), 169–176 (2006).

38. Mrosovsky, N. Masking: history, definitions, and measurement. Chronobiol. Int. 16(4), 415–429 (1999).

39. Altimus, C. M., LeGates, T. A., & Hattar, S. Circadian and Light Modulation of Behavior. in Mood and Anxiety Related Phenotypes in Mice, pp. 47–65 (Humana Press, Totowa, NJ, 2009).

40. Siepka, S. M., & Takahashi, J. S. Methods to record circadian rhythm wheel running activity in mice. Meth. Enzymol. 393, 230–239 (2005).

41. Brown, L. A., Fisk, A. S., Pothecary, C. A., & Peirson, S. N. Telling the Time with a Broken Clock: Quantifying Circadian Disruption in Animal Models. Biology. 8(1), 18 (2019).

42. Verwey, M., Robinson, B., & Amir, S. Recording and analysis of circadian rhythms in running-wheel activity in rodents. JoVE. (71), 50186 (2013)

43. Refinetti, R., Cornélissen, G., & Halberg, F. Procedures for numerical analysis of circadian rhythms. Biol. Rhythm Res. 38(4), 275–325 (2007).

44. Ruf, T. The Lomb-Scargle Periodogram in Biological Rhythm Research: Analysis of Incomplete and Unequally Spaced Time-Series, Biol. Rhythm Res. 30(2), 178–201 (1999).

45. Tackenberg, M. C., & Hughey, J. J. The risks of using the chi-square periodogram to estimate the period of biological rhythms. PLoS Comput. Biol. 17(1), e1008567 (2021).

46. Datta S., Samanta D., Sinha P., & Chakrabarti N. Gender features and estrous cycle variations of nocturnal behavior of mice after a single exposure to light at night. Physiol. Behav. 164, 113–122 (2016).

47. Jennings, M., et al. Refining rodent husbandry: the mouse: report of the Rodent Refinement Working Party. Lab. Anim. 32(3), 233–259 (1998).

